# Macrophage metallothioneins participate in the antileishmanial activity of antimonials

**DOI:** 10.1101/2020.09.30.321471

**Authors:** Deninson Alejandro Vargas, David J. Gregory, Roni Nitzan Koren, Dan Zilberstein, Ashton Trey Belew, Najib M. El-Sayed, María Adelaida Gómez

## Abstract

**Background:** Host cell functions that participate in the pharmacokinetics and pharmacodynamics (PK/PD) of pentavalent antimonials for treatment of American cutaneous leishmaniasis (CL) are critical for drug efficacy.

**Objectives:** In this study, we investigated whether macrophage mechanisms of xenobiotic detoxification contribute to drug-dependent elimination of intracellular *Leishmania*.

**Methods:** Transcriptomes of primary macrophages from CL patients (n=6), exposed ex vivo to *Leishmania* infection and Sb^V^ were generated. Candidate genes were selected and validated using short harping RNA interference (shRNA) in THP-1 cells.

**Results:** Strong induction of metallothionein (MT) genes was observed upon *Leishmania* infection and exposure to Sb^V^, with 7 MT genes (MT1 and MT2 family members) appearing within the top 20 up-regulated genes. Tandem knockdown (KD) of MT2-A and MT1-E, 1F, and 1X in THP-1 cells was achieved using a pan-MT shRNA., Intracellular parasite survival after Sb^V^ exposure was unaffected in tandem-KD cells, and this was a consequence of strong transcriptional upregulation of MTs by infection and Sb^V^, overcoming the KD effect. Gene silencing of the metal transcription factor-1 (MTF-1) abrogated expression of MT1 and MT2-A genes. Upon exposure to Sb^V^, intracellular survival of *Leishmania* in MTF-1^KD^ cells was significantly enhanced (p ≤ 0.05).

**Conclusions:** MTs are potent scavengers of heavy metals, and central elements of the mammalian cell machinery for xenobiotic detoxification. Results from this study highlight the participation of macrophage MTs in Sb-dependent parasite killing, revealing novel strategies for host-targeted optimization of antileishmanial drugs.

## INTRODUCTION

The direct activity of antimicrobials against intracellular pathogens is dependent on drug internalization into host cells and, in some instances, on host cell processes that mediate drug metabolism (activation/inactivation). This constitutes a challenge for the design of new drugs and the optimization of existing ones, because host cells constitute an additional barrier for exposure of the intracellular microbe to the drugs. Cutaneous leishmaniasis (CL) is caused by the intracellular protozoan parasite *Leishmania*, and affects more than 1.2 million people annually^1^. CL is endemic throughout Central and South America, where control remains dependent on chemotherapy with pentavalent antimonial drugs (Sb^V^). However, high rates of treatment failure (as high as 30% in controlled clinical trials), toxicity, and the difficulties associated to access to these drugs limit this control strategy^2–8^.

Exposure to Sb^V^, a metalloid closely related to arsenic, induces host stress responses that lead to activation of mechanisms of redox balance control and metal/xenobiotic detoxification^9–12^. Thus, factors mediating these stress responses could modulate Sb bioavailability within host cells, impacting the intracellular pharmacokinetics and pharmacodynamics (PK/PD) of these drugs. Sb modulates the expression of macrophage ATP Binding Cassette (ABC) transporters^13,14^. In addition, ABCC1, ABCB1 and ABCB5 have been shown to function as Sb efflux pumps in *L. donovani*^15^ and *braziliensis* infected macrophages^16^; its expression favoring intracellular parasite survival. ABCB6 can function as a plasma membrane Sb efflux transporter and as an intracellular membrane Sb importer, potentially increasing drug concentrations within the phagolysosome, favoring parasite killing^14^. These findings illustrate the participation of macrophage ABC transporters in antileishmanial drug effects.

Expression of other genes related to metal stress responses is also modulated upon Sb exposure^14,16^. Among these is metallothionein 2A (MT2-A), a small cysteine-rich cytoplasmic protein involved in zinc homeostasis and the scavenging of metals and electrophilic molecules such as reactive oxygen species (ROS) and nitric oxide (NO).^17–22^ There are four MT families members (MT1-4): MT1 and MT2 are ubiquitously expressed, whereas MT3 and MT4 are found in the central nervous system and in stratified squamous epithelium, respectively^20,23^. MTs can bind toxic metals with high affinity such as Cd, Hg, Pd, Ag, As, and Sb^19^; resulting in toxic metal tolerance and detoxification^21,23,24^. This efficient metal scavenging function results from the high thiol content of MTs and the tight regulation of MTs gene expression, which can increase more than 100 fold under metal stress, reaching intracellular concentrations of the order of millimolar^20,23,25^.

Our group and others have provided evidence of the participation of host MTs in the *Leishmania*-macrophage interactions: a) *Leishmania* infection and Sb can strongly induce the expression of MT genes in human macrophages^14,26,27^; b) an inverse correlation of MT2-A gene expression and intracellular survival of *Leishmania* during *in vitro* Sb^V^ exposure has been reported^14^; c) Sb-susceptible *L. V. panamensis* strains induce higher expression of macrophage MT2-A compared with Sb-resistant strains, suggesting strain-specific manipulation of MT2-A within macrophages^13^. Based on the above, we sought to dissect and functionally validate the participation of MTs in the antimony-mediated killing of intracellular *Leishmania*.

## MATERIALS AND METHODS

### Ethics statement

This study was approved and monitored by the institutional review board for ethical conduct of research involving human subjects of the Centro Internacional de Entrenamiento e Investigaciones Médicas - CIDEIM, in accordance with national (resolution 008430, República de Colombia, Ministry of Health, 1993) and international (Declaration of Helsinki and amendments, World Medical Association, Fortaleza, Brazil, October 2013) guidelines. All individuals voluntarily participated in the study and written informed consent was obtained from each participant.

### Reagents and chemicals

Additive-free meglumine antimoniate (MA) (Walter Reed 214975AK; lot no. BLO918690-278-1A1W601) was kindly provided by the Walter Reed Army Institute, Silver Spring, MD, USA. Phorbol-12-myristate 13-acetate was purchased from Sigma–Aldrich and zinc acetate dihydrate from J.T. Baker.

### Subjects and primary macrophages

Six adult patients, 18 to 65 years of age, with parasitological diagnosis of CL and time of lesion evolution <6 months, without apparent immune deficiencies (negative HIV test, no evidence of immunological disorder nor treatment with medication having immunomodulating effects), participated in this study. Peripheral blood samples were obtained prior to initiation of antileishmanial treatment and mononuclear cells (PBMCs) were obtained by separation using a Ficoll-Hypaque (Sigma-Aldrich) gradient.

### THP1 and primary macrophage differentiation

The human pro-monocytic cell line THP-1 and derived lines were maintained at 1 x 10^6^ cells/mL in RPMI 1640 supplemented with 10% heat inactivated FBS, 100 μg/mL streptomycin, 100 U/ml penicillin, 5 mg/mL puromycin (only for maintenance of transfected cells lines), at 37°C and 5% CO2. Monocytes were differentiated with 250 ng/mL of PMA for 3 hours, washed twice with D-PBS and cultured 24h in 6 well plates. Human PBMC-derived monocytes were differentiated to macrophages by adherence to cell culture plastic-ware as previously described^28^.

### Parasites, infection and intracellular parasite survival assays

Antimony susceptible *L. (V.) panamensis* promastigotes (MHOM/CO/2002/3594) were kept at 25°C in RPMI supplemented with 10% heat-inactivated FBS, 100 μg/mL streptomycin, 100 U/mL penicillin. Primary human macrophages and differentiated THP-1 cells were infected with human AB+ serum-opsonized stationary phase promastigotes at 10:1 *Leishmania*-macrophage ratio for 2h, washed twice with D-PBS and incubated for 24h at 34°C, 5% CO2. After infection was established, cells were exposed for 24 h or 48 h to MA (8, 16, and 32 μg/mL), or left untreated as a control. Intracellular parasite survival was measured by RT-qPCR as previously described^29^ (primers used in supplementary table S1).

### RNA isolation, cDNA library preparation, and sequence analyses

Total RNA was isolated with Trizol from uninfected, infected, and drug-treated cells. RNA quality was assessed with an Agilent 2100 Bioanalyzer using RNAnano chips (Agilent). RNA Integrity Number (RIN) ≥ 7 was considered acceptable to continue with library construction. Poly-A enriched libraries were generated using the Illumina TruSeq Standard mRNA preparation kit and checked via the Bioanalyzer and quantitative PCR (KAPA Biosystems). Paired-end reads (100 nt) were obtained using the Illumina HiSeq 1500 (BioProject ID PRJNA633893 and supplementary table S2). Fastqc^30^ was used to evaluate sequencing quality; Trimmomatic^31^ filtered low-quality reads and trimmed bases when the mean quality score fell below a threshold phred score of 20. Reads were mapped against the human^32^ (hg38) and *L. V. panamensis*^33^ genomes (v36ish) using tophat^34^. HTSeq^35^ was used to count reads mapping to each gene feature. The count tables were restricted to the set of protein coding genes and filtered to remove non-expressed and very weakly expressed genes. The remaining genes were assessed for significant outliers and batch effects by visualizations of normalized data. Library sizes and count densities were calculated on non-normalized data; pairwise correlation and distances, outlier detection, and principal component analysis (PCA) was performed on log2, cpm, quantile normalized data with and without accounting for batch in the model or surrogate estimation with sva (using svaseq or combat)^36^. DESeq2^37^ was used to perform differential expression analyses alongside a statistically uninformed basic method as a negative control (supplementary table S3). Differentially expressed genes were contrasted between control vs. infected and MA treated primary human macrophages. Genes deemed significantly different according to DESeq2 (| logFC | > 1.0 and a FDR adjusted p-value < 0.05) were passed to various ontology tools. Enrichment PPI network analysis was carried out using STRING 10.

### Short hairpin RNA (shRNA) constructs

A lentivirus-based system was used for shRNA-mediated gene silencing in THP-1 monocytes as previously described^38^. At least two independent sets of oligonucleotide pairs for gene knockdown of human MTs and the transcription factor MTF-1 (supplementary table S4) were synthesized and cloned into the pLKO.1-TCR vector (Addgene, Cambridge, MA, USA); the same vector was also used as an empty vector control. Lentiviral particles were generated by co-transfection of endotoxin-free hairpin-containing pLKO.1-TCR, psPAX2 and MD2.G (Addgene) into HEK-293T cells. FuGENE HD (Roche) was used as the transfection reagent. Lentivirus-containing cell supernatant was collected 4 days after transfection and subsequently used to transduce THP-1 monocytes in medium containing 10 mg/mL polybrene in a proportion 1:1. Transduced cells were selected under puromycin pressure (5 mg/mL) for a minimum of 5 days. Gene knockdown was confirmed by RT–qPCR using SYBR green (Applied Biosystems) and TaqMan^®^ Gene Expression Assays (Applied Biosystems, supplementary table 2). shRNA transduction was confirmed by DNA sequencing.

### Cytotoxicity assays

PMA-differentiated THP-1 cells and derived cell lines were exposed to a dose range (8 μg/mL – 256 μg/mL) of MA for 72 hours. Cell viability was assessed by MTT assay (ATCC) (supplementary figure S1).

#### Statistical analysis

Based on the distribution of the data, differences in gene expression were tested with one-way analysis of variance and Tukey’s multiple comparisons test. Differences in variance for the remaining experiments were analyzed with unpaired *t* test. A significance level of *p* ≤ 0.05 was used for all statistical tests. Statistical analyses were performed using GraphPad Prism software (version 6).

## RESULTS

### Metallothioneins are within the top 20 transcripts up-regulated by *Leishmania* infection and exposure to Sb^V^ in macrophages from CL patients

Peripheral blood mononuclear cell (PBMC)-derived macrophages from CL patients (n=6) were infected *ex-vivo* with *L.V. panamensis* and exposed to Sb^V^ (32μg-Sb/mL, as meglumine antimoniate - MA). Following RNAseq data collection, a total of 16,841 transcripts were detected. Principal Component Analysis (PCA) showed separation between uninfected/untreated control macrophages and those infected with *L.V.panamensis* and exposed to MA (Figure 1A). After filtering of the differential expression (DE) data by a |logFC| ≥ 2 and p ≤ 0.05, a set of 106 transcripts remained, of which 69 were up-regulated and 37 down-regulated (Supplementary table 1). Interestingly, among the top twenty up-regulated transcripts, seven were metallothionein genes: MT1-E, MT1-F MT1-M, MT1-G, MT1-H, MT1-X and MT2-A (Figure 1B and Supplementary table 1).

**Figure 1.**
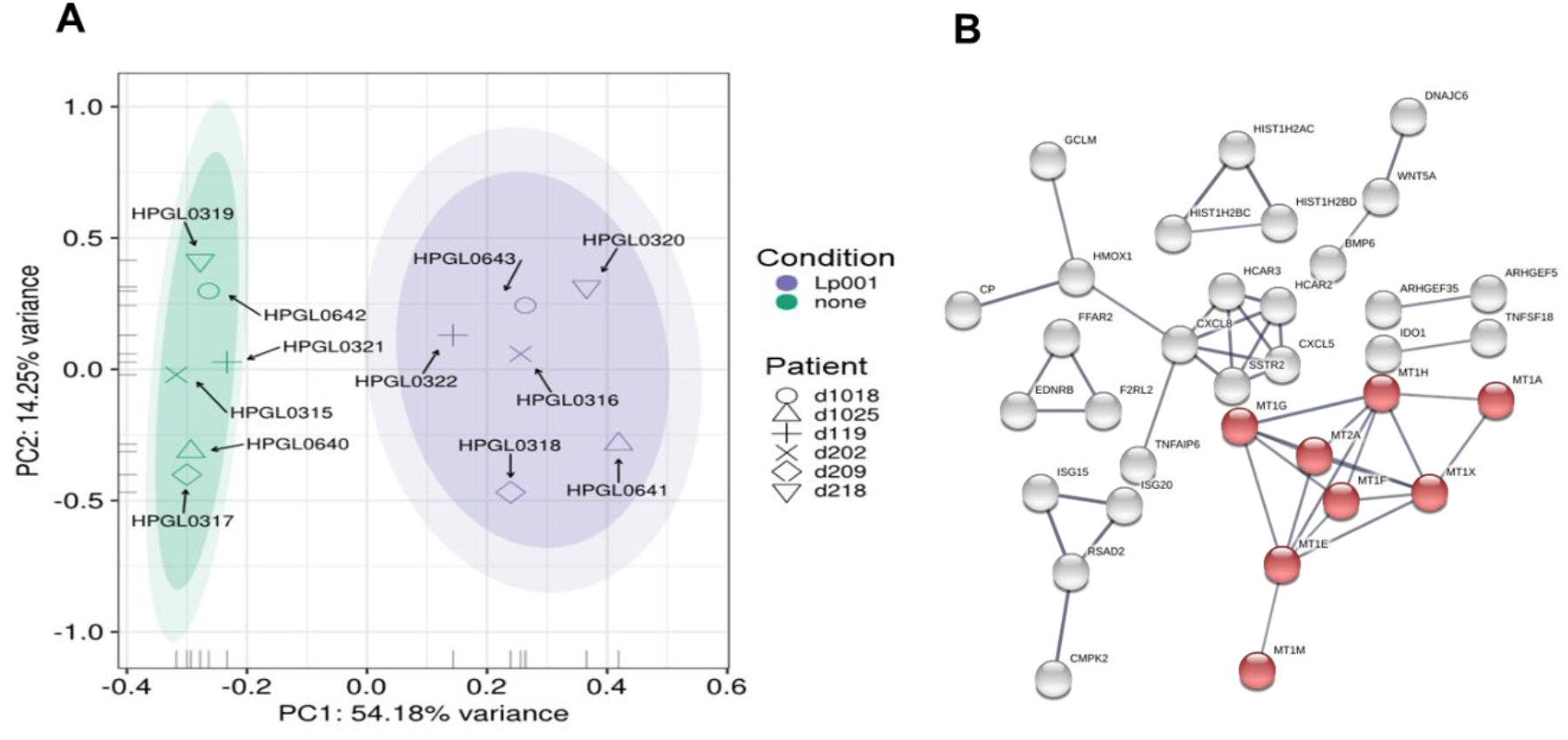
PCA plot and network analysis of macrophage transcriptomes. A) PCA plot of RNAseq data after batch correction (each donor as an independent batch), of PBMC-derived macrophages from CL patients (n=6), which were infected *ex-vivo* with *L.V. panamensis* (Lp001) and exposed to MA (32μg-Sb/mL) for 24 h (purple), or were left uninfected and untreated as controls (green). Ovals represents confidence Interval-CI: 90% (inner) and 95% (outer). Symbols represent samples from each patient. B) STRING network analysis of up-regulated genes with a |logFC| cutoff ≥ 2. Nodes represent proteins and edges represent protein-protein interactions (PPI) with the confidence of interaction set at 0.7. Highlighted in red is the metallothioneins PPI network.

### Induction of MTs expression by zinc acetate enhances MA-mediated killing of *Leishmania*

To discern how MTs participate in the activity of antimonials, THP-1 cells were exposed to Sb^V^ and the expression of MT1-E, 1-F, 1-X, MT2-A, MT3 and MT4 was quantified (primer sequences available in Supplementary table 2). Expression of MT1-X was the highest, peaking at 8-fold induction over untreated cells; followed by MT1-E, MT2-A and MT1-X (Figure 2A). MT3 and MT4 transcripts were not detected in macrophages, consistent with their tissue specificity. Next, we explored whether maximal induction of MTs in THP-1 cells would enhance the Sb^V^ mediated killing of *Leishmania*. To do so, THP-1 cells were exposed to a dose range of 25 μM to 400 μM Zn acetate for 24h, and peak expression of MTs (>100 fold) was observed with 400 μM Zn acetate^23^ (Figure 2B). Therefore, THP-1 cells were pre-treated for 24h with 400 μM Zn acetate, followed by *L. V. panamensis* infection for additional 24h, and exposed to increasing and non-cytotoxic concentrations of MA (Supplementary figure 1). Pre-treatment with Zn acetate increased over 40% the Sb-dependent intracellular elimination of *L.V. panamensis* (Figure 2C), supporting the contribution of MTs to this phenotype.

**Figure 2.**
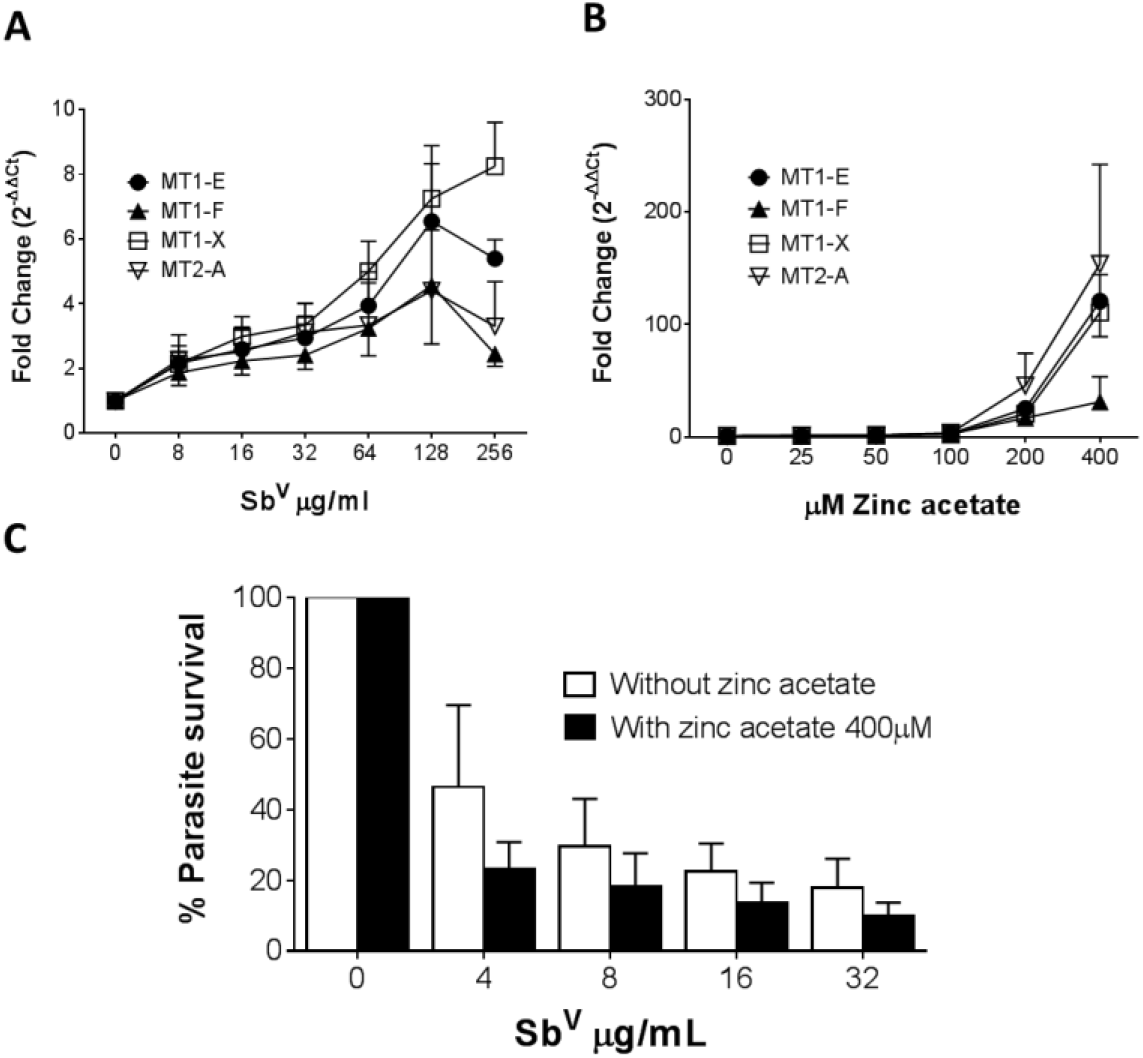
MTs are induced by zinc acetate and antimony. **A)** Fold change gene expression of MTs in THP-1 cells exposed to sub-cytotoxic doses of Sb^V^. **B)** Fold change gene expression of MT1-E, 1-F, 1-X and MT2-A in THP-1 cells exposed to increasing doses of zinc acetate. **C)** *L.V. panamensis* survival (%) in THP-1 cells pre-treated with 400 μM Zn-acetate for 24h and exposed to Sb^V^ (8, 16 and 32 μg/mL). THP-1 gene expression and parasite survival were quantified by RT-qPCR. Each experiment was run as 3 independent replicates. Data are shown as mean ± SD.

### Strong transcriptional up-regulation of MTs abrogates shRNA silencing of MT genes

Taking advantage of the high sequence similarity of MT genes, a shRNA was constructed which targets the tandem knockdown (MT_tandem^KD^) of MT1 and MT2 family member genes (Figure 3A and Supplementary table 3). Expression of MT1-E, 1F, 1X and MT2-A was efficiently silenced as shown in Figure 3B. To explore the phenotypic effects of MTs knockdown on intracellular survival of *Leishmania*, MT_tandem^KD^ cells were infected with *L. V. panamensis* and exposed to Sb^V^. Despite efficient knockdown of MT genes, intracellular parasite survival remained unchanged in MT_tandem^KD^ cells compared to empty-vector transfected control cells (Figure 3C). Considering that expression of MT genes is strongly induced by *Leishmania* and Sb^V^, we questioned whether MTs knockdown was maintained during the experimental conditions (infection and drug exposure). Despite the efficient tandem knockdown of MTs at basal conditions, their strong transcriptional up-regulation during *Leishmania* and Sb^V^ exposure overcame the shRNA-silencing effect (Figure 3D); partially explaining why parasite survival was similar in MT_tandem^KD^ and empty-vector control cells.

**Figure 3.**
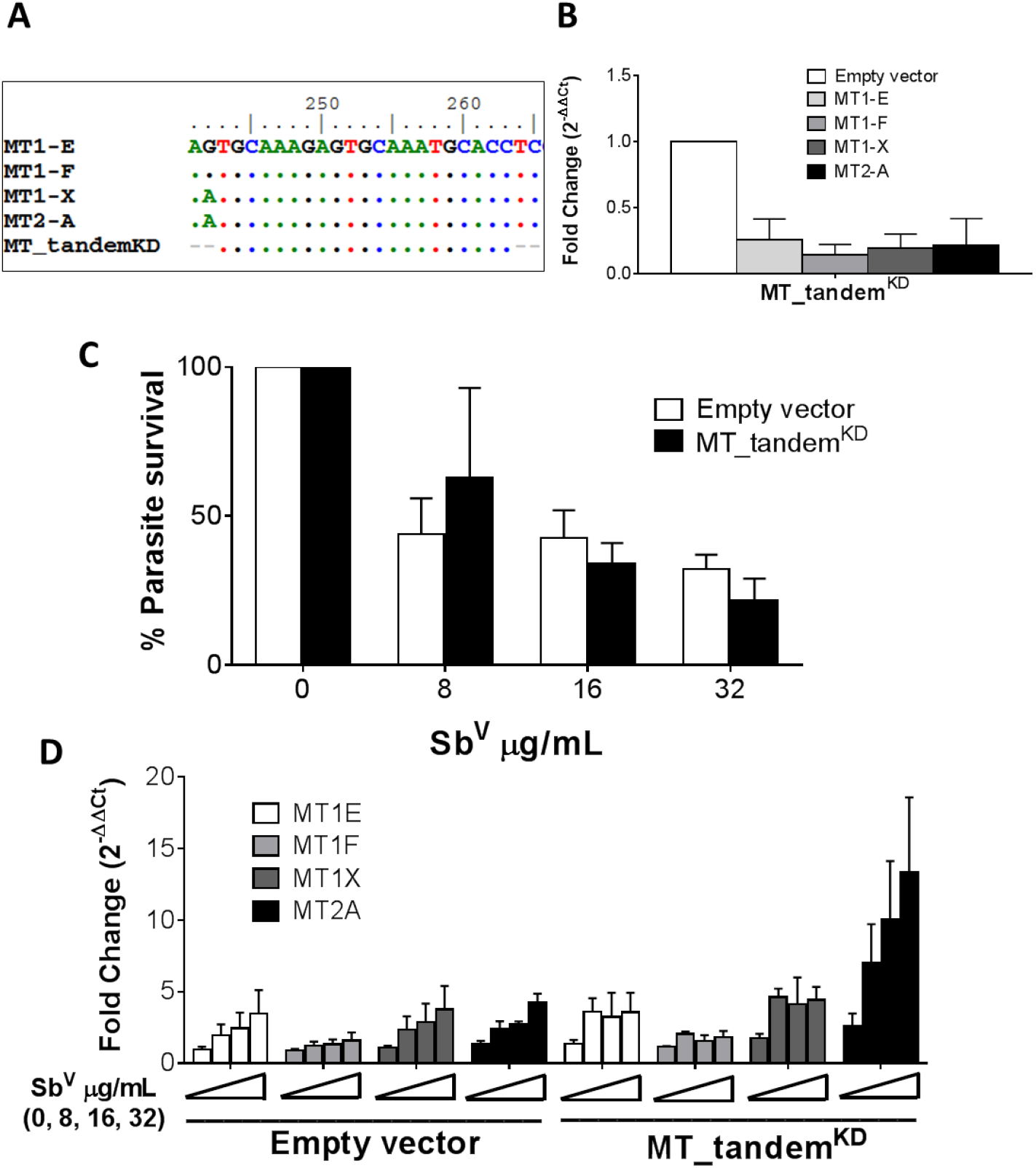
MTs tandem knockdown is abrogated by strong transcriptional up-regulation. **A)** Sequence alignment of MT genes and MT_tandem^KD^ oligo sequence showing the conserved region targeted for tandem shRNA. **B)** Validation of the tandem KD shown by MTs gene expression in uninfected MT_tandem^KD^ and empty-vector control cells. **C)** Percentage of parasite survival in empty vector control and MT_tandem^KD^ cells infected with *L.V. panamensis* and exposed to Sb^V^ (8, 16 and 32 μg/mL). **D)** Fold change gene expression of MTs upon infection and exposure to Sb^V^ (8, 16 and 32 μg/mL) in empty vector control and MT_tandem^KD^ cells. Gene expression and parasite survival were quantified by RT-qPCR. Each experiment was performed in 3 independent replicates. Data are presented as mean values ± SD.

### Expression of MTs is silenced by knockdown of MTF-1 and favors survival of intracellular *Leishmania*

Metal transcription factor-1 (MTF-1) is the principal transcription factor regulating expression of MTs genes during metal-induced stress.^39^ We explored whether knockdown of MTF-1 could limit the transcriptional induction of MT genes in our experimental conditions. Using shRNA, the steady-state level of MTF-1 was knocked down by 50% (Figure 4A). Expression of MTs in MTF-1^KD^ cells was evaluated under basal conditions and upon infection with *L.V. panamensis* and exposure to Sb^V^. A slight reduction (ranging from 5% to 40%) of MT1 and MT2 genes expression was observed in uninfected and unstimulated MTF-1^KD^ cells (Figure 4B). However, induction of MT1-E, F, X and MT2-A expression was completely repressed in *L.V. panamensis* infected cells subsequently exposed to Sb (Figure 4C).

**Figure 4.**
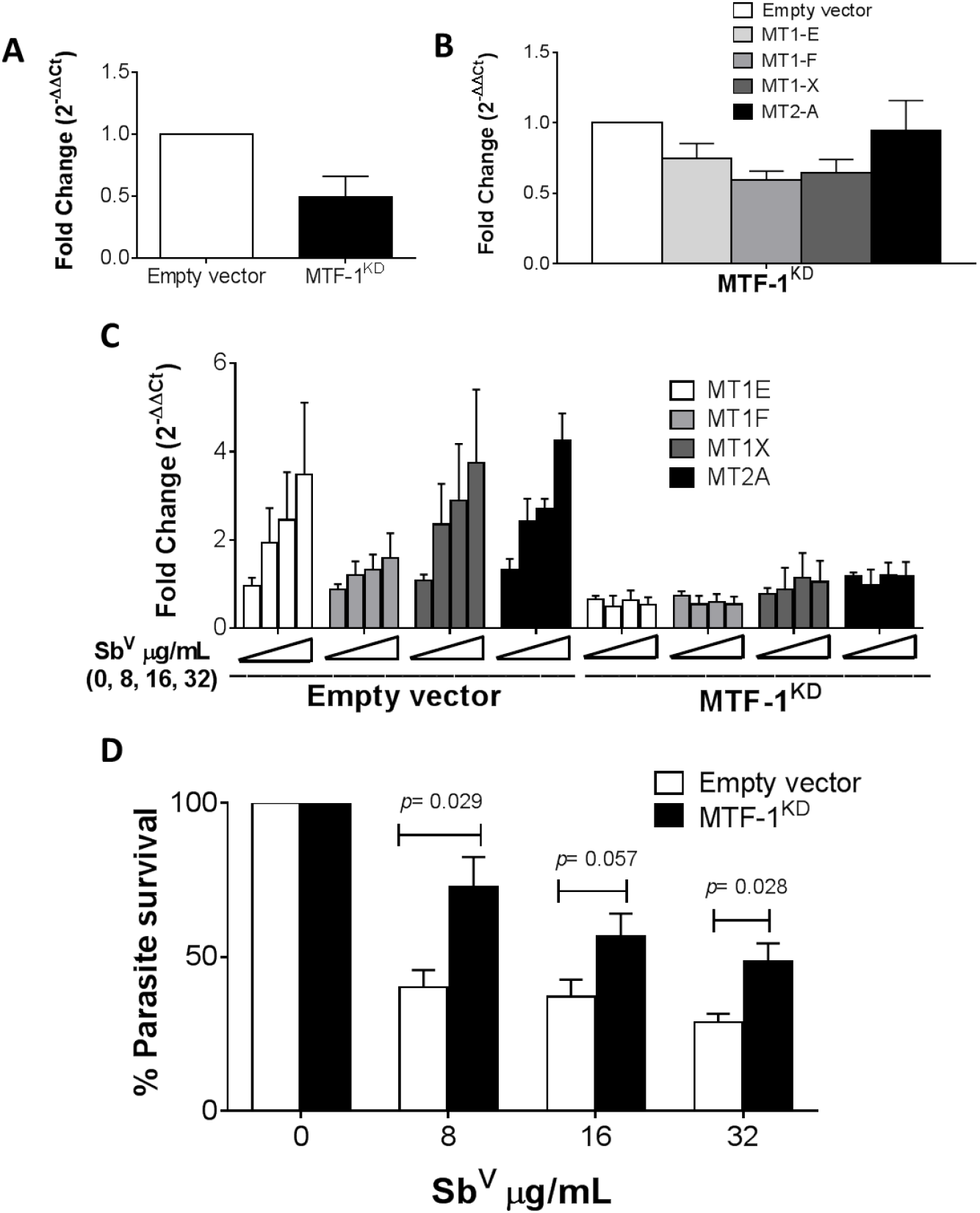
Induction of MTs is abolished by knockdown of MTF-1 favoring intracellular parasite survival. **A**) Fold change gene expression of MTF-1 in empty vector control and MTF-1^KD^ THP-1 cells. **B**) Fold change gene expression of MTs in unstimulated or **C)** infected and Sb^V^ (8, 16 and 32 μg/mL) exposed MTF-1^KD^ cells. **D**) Percentage of intracellular *L.V. panamensis* survival after Sb^V^ exposure. Gene expression and parasite survival were quantified by RT-qPCR. Each experiment was run in 3 independent replicates. Mann-Whitney test was used for statistical analysis. Data are presented as mean values ± SD.

MTF-1^KD^ and empty vector control THP-1 cells were infected and exposed to Sb^V^ (8 - 32 μg/mL), and intracellular parasite survival measured by RT-qPCR. Sb-dependent parasite killing was efficient in empty-vector control cells, where parasite survival was below 50% for all Sb^V^ doses tested (Figure 4D). In contrast, *Leishmania* survival after drug exposure significantly increased in MTF-1^KD^ cells compared to empty vector control cells (Figure 4D), and was above 50% for all evaluated doses and up to 75% in the 8 μg/mL Sb^V^ dose. These data suggest that MTF-1^KD^ favors intracellular survival of *Leishmania* by impairing MTs gene expression.

## DISCUSSION

Metallothioneins were initially reported in the early ‘70s and characterized as cadmium binding proteins with a role in metal detoxification^40^. Subsequent studies demonstrated that these proteins could also bind other metals including Sb^19^, leading to protection against metal-induced toxicity. Pentavalent antimonials continue to be first line treatment for leishmaniasis in many endemic countries^41^. Despite more than 100 years of use, the mechanisms of action and of exposure of the intracellular parasite to the active drug (Sb^III^), remain poorly understood. Recently, the role of host cells in antimony metabolism and detoxification has gained increased attention due to their potential participation in treatment outcome, and as potential targets for optimization of drug exposure^13,14,16^. Here, we provide evidence of the participation of macrophage metallothioneins in the Sb-mediated killing of intracellular *Leishmania*.

Studies in MT1/MT2-null mice have demonstrated increased toxicity of cisplatinum, Cd, Hg, Cu, Zn and As -a metalloid closely related to Sb-^21,24,42–44^. Concurring with these findings, tolerance to metal-induced hepato- and nephrotoxicity has been demonstrated in transgenic mice overexpressing MTs^40,45,46^. Although the precise mechanism by which MTs facilitate Sb-dependent killing of *Leishmania* remains to be determined, the metal scavenging function of MTs could promote Sb accumulation within infected macrophages. Interestingly, it has been shown that MTs translocate to lysosomes^40,47,48^, suggesting that MT-Sb complexes could be found within lysosomes. *Leishmania* resides in phagolysosomal compartments within host cells. Therefore, as a consequence of the biological process of phagosome-to-phagolysosome maturation, fusion of MT-Sb containing lysosomes with *Leishmania*-containing phagosomes could result in enhanced exposure of the intracellular parasite to the drug.

Pentavalent antimonials are pro-drugs which need to be reduced to the trivalent active form to exert their antileishmanial activity (Sb^V^→Sb^III^). MTs have an important function in the cellular redox balance due to their high thiol content^17–22^. Under stress conditions, the strong up-regulation of MTs gene expression results in a redox capacity that can surpass that of GSH^20,23,25^. High GSH content has been shown to promote Sb^V^ to Sb^III^ reduction in mammalian cells as well as in *Leishmania*^49,50^. Thus, MTs could participate in the reduction Sb^V^ to Sb^III^ favoring parasite elimination.

MTF-1 is the main transcription factor involved in MTs expression during metal stress responses, and partially during oxidative stress exposure/response^22,51–54^. Our results and those of others demonstrate that MTF-1 knockout/knockdown efficiently abolishes cellular expression of MTs under metal and oxidative stress ^22, 39, 53^. Although our data provide evidence that repression of MTs expression via MTF-1 gene knockdown promotes intracellular survival of *Leishmania* after exposure to Sb, we cannot rule out the intervention of other MTF-1 mediated mechanisms in the enhanced parasite survival, such as those involved in Zn transport^55–59^.

Our results support a dual role of MTs in toxic as well as therapeutic metal binding, highlighting the potential to harness host cell redox and metal detoxification systems to enhance drug bioavailability and exposure targeted to intracellular pathogens. These findings enlighten interesting drug-related homeostatic processes occurring during treatment of intracellular microbe infections, whereby the same mechanism that promotes host protection to drug-induced toxicity, in this case against Sb-induced stress, can enhance the antimicrobial activity of the drug.

## ACKNOWLEDGEMENTS

We gratefully acknowledge the patients and volunteers who participated in this study and the members of the Clinical Unit of CIDEIM in Cali and Tumaco for recruitment of participants and follow-up. This work was conducted in partial fulfillment of the requirements for the DSc degree in Biomedical Sciences of Universidad del Valle to DAV.

## FUNDING

This work was supported in part by US National Institutes of Health (NIH) Grant R01AI104823 (https://www.niaid.nih.gov/) and Wellcome Trust award 107595/Z/15/Z to MAG. DAV was supported by COLCIENCIAS DSc student award 647.

## TRANSPARENCY DECLARATIONS

The authors declare that they have no conflicts of interest with the contents of this article. The content is solely the responsibility of the authors and does not necessarily represent the official views of the National Institutes of Health. The funders had no role in study design, data collection and analysis, decision to publish, or preparation of the manuscript.

## Notes

### Competing Interest Statement

The authors have declared no competing interest.

